# Morphological and transcriptomic evidence for ammonium induction of sexual reproduction in *Thalassiosira pseudonana* and other centric diatoms

**DOI:** 10.1101/144667

**Authors:** Eric R. Moore, Briana S. Bullington, Alexandra J. Weisberg, Yuan Jiang, Jeff Chang, Kimberly H. Halsey

## Abstract

The reproductive strategy of diatoms includes asexual and sexual phases, but in many species, including the model centric diatom *Thalassiosira pseudonana*, sexual reproduction has never been observed. Furthermore, the environmental factors that trigger sexual reproduction in diatoms are not understood. Although genome sequences of a few diatoms are available, little is known about the molecular basis for sexual reproduction. Here we show that ammonium reliably induces the key sexual morphologies, including oogonia, auxospores, and spermatogonia, in two strains of *T. pseudonana*, *T. weissflogii*, and *Cyclotella cryptica*. RNA sequencing revealed 1,274 genes whose expression patterns changed when *T. pseudonana* was induced into sexual reproduction by ammonium. Some of the induced genes are linked to meiosis or encode flagellar structures of heterokont and cryptophyte algae. The identification of ammonium as an environmental trigger suggests an unexpected link between diatom bloom dynamics and strategies for enhancing population genetic diversity.

## Introduction

Diatoms are protists that form massive annual spring and fall blooms in aquatic environments and are estimated to be responsible for about half of photosynthesis in the global oceans [1]. This predictable annual bloom dynamic fuels higher trophic levels and initiates delivery of carbon into the deep ocean biome. Diatoms have complex life history strategies that are presumed to have contributed to their rapid genetic diversification into ~200,000 species [2] that are distributed between the two major diatom groups: centrics and pennates [3]. A defining characteristic of all diatoms is their restrictive and bipartite silica cell wall that causes them to progressively shrink during asexual cell division. At a critically small cell size and under certain conditions, auxosporulation restitutes cell size and prevents clonal death [4-6]. The entire lifecycles of only a few diatoms have been described and rarely have sexual events been captured in the environment [7-9].

So far, all centric diatoms appear to share the process of oogamous sexual reproduction (Fig 1). The average cell size of a population of asexually dividing diatoms decreases as a result of differential thecae inheritance. At a critically small size, cells become eligible to differentiate into male and female cells. Meiosis in the male spermatogonangium produces multinucleate spermatogonia that divide into individual haploid spermatocytes. Meiosis in the female oogonia produces a single functional haploid nucleus that is fertilized by a flagellated spermatocyte through an opening in the oogonia thecae. Fertilized oogonia expand into a large auxospore where new, large thecae are formed for the new, enlarged initial cell. Auxosporulation can also occur asexually, but it is considered an ancillary pathway for cell size restitution in diatom species that have a sexual path for reproduction [5].

**Fig 1.**
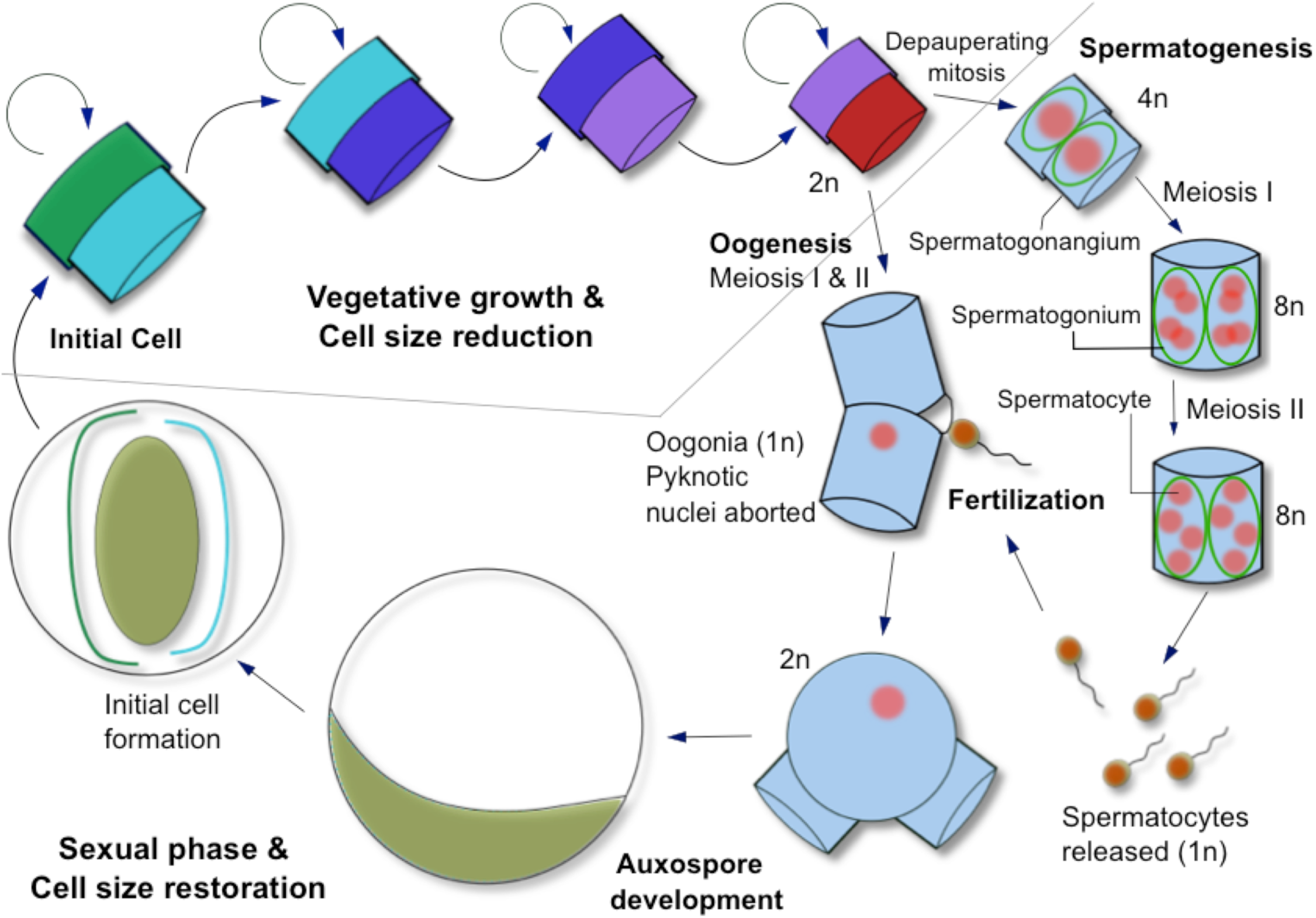
The life cycle of a centric diatom. The average cell size of a population of asexually dividing diatoms decreases as a result of differential thecae inheritance. At a critically small size, cells can initiate sexual reproduction and differentiate into male and female cells. Meiosis in the male spermatogonangium produces multinucleate spermatogonia that divide into individual haploid spermatocytes. Meiosis in the female oogonia produces a single functional haploid nucleus that is fertilized by a flagellated spermatocyte through an opening in the oogonia thecae. Fertilized oogonia expand into a large auxospore where new, large thecae are formed for the new initial cell.

The environmental factors that trigger formation of sexual cells and sexual reproduction in centric diatoms are not well understood [ 10, 11], but sexualization appears to be strongly associated with conditions causing synchronous sexuality in cells experiencing growth stress [12]. Besides the size threshold requirement, previous observations indicate that sexualization is possible when active growth has ceased, causing cell cycle arrest [13, 14] and cell densities are sufficient to permit successful fertilization of the oogonia by the spermatocyte [15]. Light interruption with an extended dark period [13], changing salinities, and nutrient shifts [16], have sometimes been successful in inducing sexual reproduction, probably by causing cell cycle arrest. Recently, pheromones produced by the pennate diatom, *Seminavis robusta*, have been identified that cause cell cycle arrest and induce the sexual pathway [17]. However, we are aware of no method that reliably causes induction of all of the sexual stages of centric diatoms shown in figure 1.

The ecological importance of diatoms, combined with their potential uses in materials chemistry, drug delivery, biosensing [18, 19], and bioenergy [20, 21], prompted genome sequencing of *T. pseudonana* CCMP1335 (a ‘centric’ diatom collected from the North Atlantic Ocean) and *Phaeodactulum tricornutum* (a ‘pennate’ diatom), which have become model organisms for experimental studies [22, 23]. However, sexual morphologies have never been observed in either of these species or in the vast majority of diatoms [10]. The inability to reliably control the sexual cycle in centric diatoms has severely hindered studies to understand the silica deposition process, as well as the genetic regulation, ecology, and evolution of sex [10, 24, 25]. Both of the model diatoms were thought to have repurposed their extant genetic toolkits and lost the need and ability for a sexual lifestyle [10, 11, 26].

Here we show that two strains of *T. pseudonana* and two other centric species, *T. weisflogii* and *Cyclotella cryptica*, can be reliably induced into the sexual reproductive pathway when cells are below the critical size threshold and exposed to ammonium during the stationary phase of growth. Ammonium induced oogonia, auxospore, and spermatocyte formation in each of these species. Induction of sexuality was further supported by RNA sequencing (RNAseq) which revealed 1,274 genes whose expression patterns changed when *T. pseudonana* became sexualized by ammonium. Meiosis genes and genes associated with flagellar structures of heterokont and cryptophyte algae were differentially expressed in ammonium-induced cells compared to nitrate grown cells. We anticipate that this discovery will open opportunities to study the evolution of diatom lifecycles and facilitate expansion of diatom breeding to explore functional genetics for molecular ecology, nanotechnology and biofuels applications.

## Results and discussion

### Ammonium triggers sexual morphologies

We observed *T. pseudonana* CCMP1335 cell morphologies consistent with sexual reproduction when cells were propagated in artificial seawater medium supplemented with ammonium. The proportion of cells that differentiated into sexual cell types was dependent on ammonium concentration, with up to 39% of the population identified as oogonia or auxospores in cultures supplemented with 800 μM NH_4_Cl (Fig 2A). Oogonia and auxospores were first observed at the onset of stationary phase and reached maximum population proportions in late stationary phase (Fig 2A). Ammonium also induced oogonia and auxospore production in *T. pseudonana* CCMP1015 (collected from the North Pacific Ocean), *T. weissflogii*, and *Cyclotella cryptica* (S1 Fig). A few oogonia and auxospores were observed in nitrate grown cultures with no added ammonium (Fig 2A and S1). However, with the exception of nitrate grown *C. cryptica* cultures, which generated oogonia and auxospores constituting 11% of the total population, oogonia and auxospores were only a small percentage of the total population in nitrate-grown *T. pseudonana* and *T. weissflogii.* Even though auxospores can have diameters 3-4 times that of asexual cells, such small population proportions do not lead to discernable shifts in cell size distributions obtained by particle size analysis (e.g., Coulter counter, a commonly used method to assess population size). We initially observed auxospores when performing visual inspections using a light microscope of our cultures that were growing in ammonium. For the data reported here, oogonia and auxospores were quantified by manually counting the cell types using a hemocytometer. We suspect that reliance on laboratory instruments such as particle counters and flow cytometers in place of microscopic analysis is one reason that sexual morphologies in these well-studied diatom species have gone undetected until now. Laboratory stock cultures are typically maintained in media with low concentrations of nitrogen, especially when ammonium is supplied as the nitrogen source because it has been considered to be toxic to diatoms in high concentrations [27]. Therefore, it may not be surprising that sexual cells have gone un-noticed due to the low rates of sexual induction in the presence of low ammonium concentrations (Fig. 2A).

**Fig 2.**
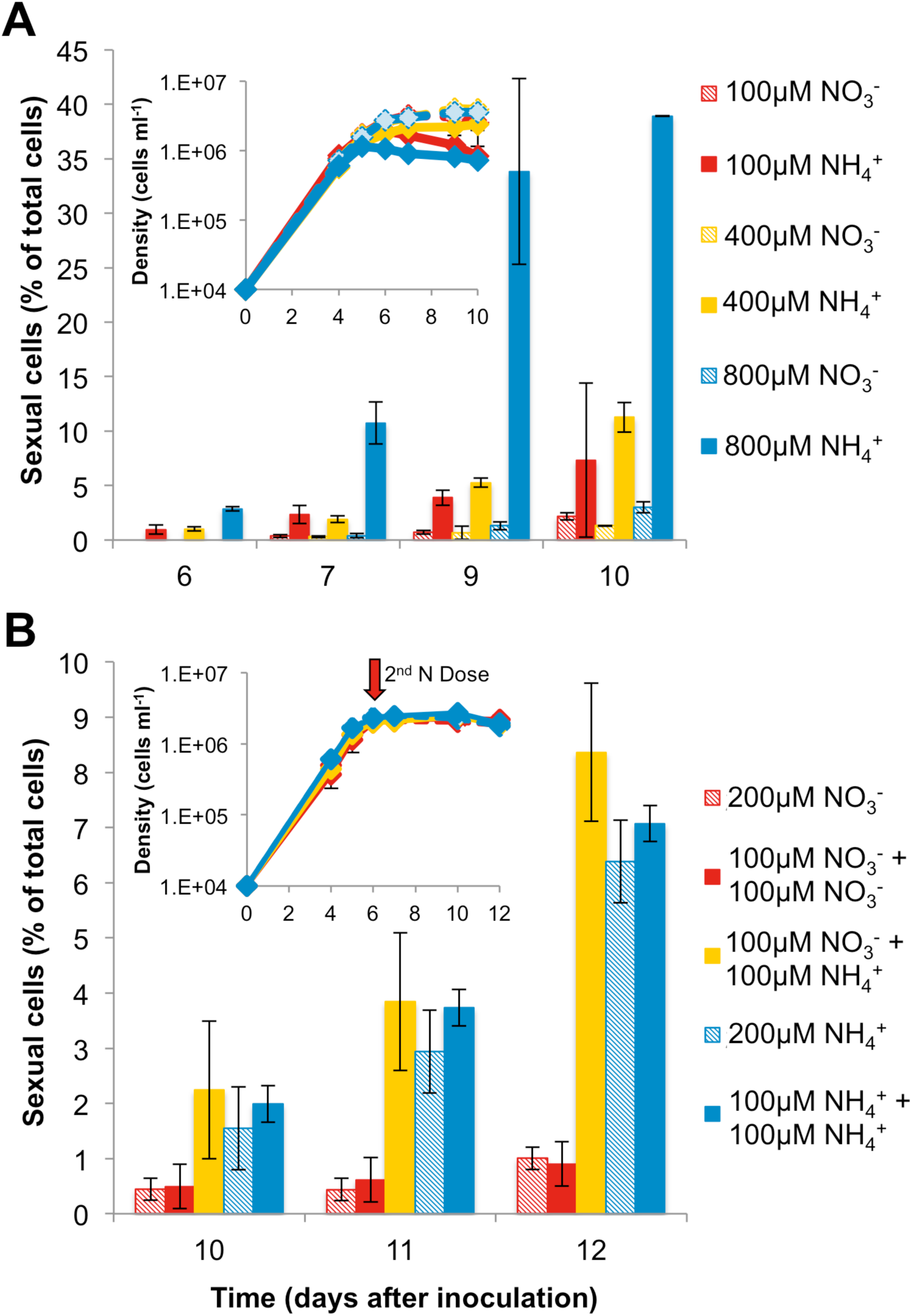
Ammonium induces sexual morphologies in *T. pseudonana* CCMP1335. **(A)** Proportion of sexual cells (oogonia and auxospores) relative to the total population in cultures of *T. pseudonana* grown in the presence of NH_4_Cl or NaNO_3_; n=3 independent cultures, average of 300 cells counted per replicate. Oogonia and auxospores were only observed beginning in stationary phase, data are mean values, error bars are s.d.. Inset: corresponding growth curve linking the onset of stationary phase with first appearance of sexual cells on day six. (**B)** Sexual cells were observed in cultures with NH_4_Cl present at inoculation (blue hatched and solid blue bars) or following NH_4_Cl addition at the onset of stationary phase (yellow bars). Legend shows concentration of nitrogen source provided at inoculation and concentration of nitrogen source added at the time of the second dosing. Two control treatments were supplied 200 μM nitrogen source at inoculation only. Inset: corresponding growth curve showing the onset stationary phase and timing of 2^nd^ nitrogen addition; n=3 independent cultures, average of 281 cells counted per replicate, data are mean values, error bars are s.d..

Cells differentiated into oogonia and auxospores regardless of whether ammonium was supplied at inoculation or at the onset of stationary phase (Fig. 1B). Thus, it appears that stationary phase and ammonium availability are key factors that trigger formation of sexual cells in centric diatoms. Resource depletion can arrest the cell cycle [ 14], and the presence of ammonium at the onset of stationary phase appears to activate the sexual cycle. Auxospore formation was observed in *Cyclotella meneghiniana* during stationary phase [28], and other protists initiate sex under stress in response to nutrient depletion or oxidative DNA damage [29]. Ammonium can inhibit photosynthesis [27]; however, diatoms, including *T. pseudonana*, can acclimate to millimolar ammonium concentrations [30]. It is possible that high ammonium concentrations intensify the stress condition required for the sexual pathway. Nevertheless, ammonium consistently caused formation of ten-fold more sexualized cells than the same concentrations of nitrate (Fig 2 and S1).

Our results showing that ammonium induced formation of sexual cells in several centric diatom species suggests that it may serve as a key environmental factor regulating the sexual lifecycle across centric diatoms. Ammonium is typically present in very low concentrations in aquatic ecosystems. However, ammonium reached 12.6 mM in a eutrophic lake where the centric diatom, *Aulacoseira subarctica*, was observed undergoing sexual reproduction [8]. Clearly, ammonium was not the growth-limiting nutrient under those conditions or in our laboratory cultures (Fig 2). *Pseudo-nitzschia* auxospore formation was positively correlated with ammonium, which was measured to be 14 μM during a major bloom event off the coast of Washington [7]. Thus, the formation of sexual cells appears to be triggered by the presence of ammonium while at least one other growth factor becomes limiting, such as light (discussed below), phosphorous, silica [7], vitamins, or trace elements.

Cell differentiation in *T. pseudonana* was induced irrespective of growth rate in exponential phase, light intensity, or light regime. However, the growth parameters did affect the proportion of differentiated cells. Oogonia and auxospores were only 0.5% of the population when grown under very low light (5 μE) with 200 μM NH_4_Cl and increased to 39% when grown under moderate light (100 μE) and 800 μM NH_4_Cl (Table 1). The proportions of oogonia and auxospores increased with light up to moderate intensities (70-100 μE) but decreased at high (220 μE) intensities (Table 1), suggesting that photon flux has an important role in meeting the energetic demands of sexual reproduction. Other work showed sexualization was more prevalent at light <50 μE [31] or with the addition of a dark period [13, 32, 33]. Likely, the optimum light intensity or need for a dark period to precede sexual induction [34] is species-specific and linked to adaptive life histories [5].

**Table 1.** Effects of growth parameters on induction of sexual reproduction in *T. pseudonana* CCMP1335, *T. weissflogii* and *C. cryptica*. Oogonia and auxospores always appeared in stationary phase. The percentage of the total population (at least 300 cells counted per replicate) differentiated into oogonia or auxospores when grown in nitrate or ammonium is shown for the day they were at their maximum number; data are mean values ± s.d., biological replicates n=3.

| Species | Light intensity (μE) | Light regime | Nitrogen concentration (μM) | Specific growth rate ($\text{d}^{-1}$) | Oogonia and auxospores in $\text{NO}_3^-$ (%) | Oogonia and auxospores in $\text{NH}_4^+$ (%) |
| --- | --- | --- | --- | --- | --- | --- |
| *T. pseudonana* | 5 | 24 h | 200 | 0.28 | 0.21 ± 0.22 | 0.52 ± 0.52 |
| | 50 | 24 h | 200 | 0.89 | 3.67 ± 0.55 | 7.67 ± 0.50 |
| | 50 | 12 h/12 h L/D | 200 | 0.80 | 0.67 ± 0.27 | 2.32 ± 0.52 |
| | 220 | 24 h | 200 | 1.2 | 0.26 ± 0.26 | 1.30 ± 0.91 |
| | 70 | 24 h | 800 | 0.98 | 3.01 ± 0.52 | 38.9 ± 0.04 |
| *C. cryptica* | 100 | 24 h | 800 | 0.38 | 11.1 ± 2.27 | 37.6 ± 3.71 |
| *T. weissflogii* | 100 | 24 h | 800 | 0.52 | 4.50 ± 2.36 | 39.8 ± 5.65 |

### Visualization of sexual morphologies

Confocal, light, and scanning electron microscopy were used to document the cell morphologies at various stages in the life history cycle of *T. pseudonana* (Fig 3 and S2). Oogonia were elongated relative to asexually growing vegetative cells and exhibited a bent morphology and swelling of the plasma membrane at the junction of the hypotheca and epitheca (Figs 3B, 3C and S2B-D). In oogonia, cellular contents became localized to the ends of the cell resulting in an apparent empty space near the area of membrane swelling where fertilization likely occurs [5].

**Fig 3.**
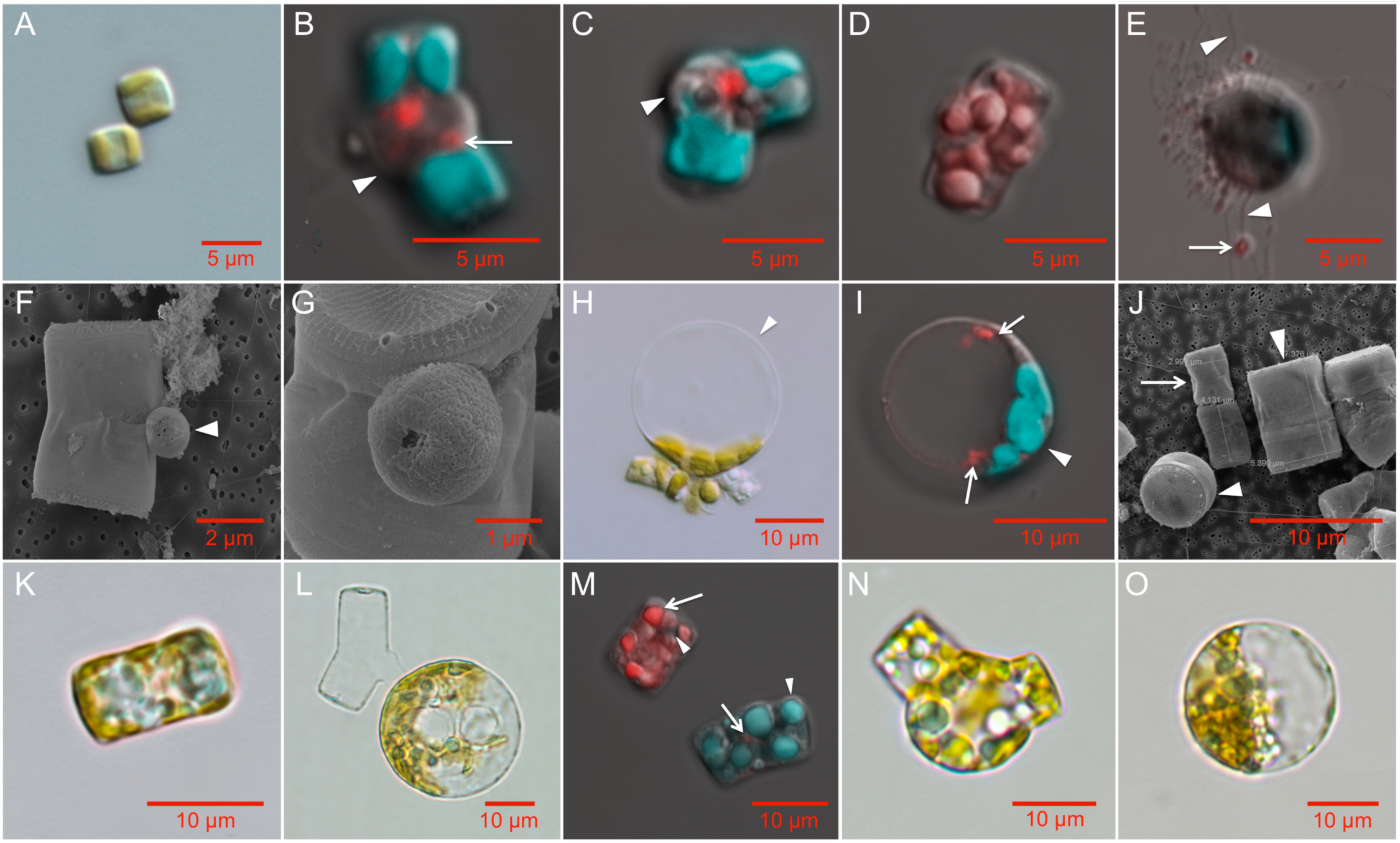
The life cycle stages of *T. pseudonana* (A-K), T. weissflogii (L) and C. cryptica (M-O) imaged using scanning electron microscopy (SEM), light (LM), and confocal microscopy (CFM). **A:** Two vegetative cells (LM, CCMP1335). **B:** Oogonium displaying separation of thecae (arrowhead) and putative pycnotic nucleus indicated by the arrow (CFM, CCMP1015). **C:** Oogonium sharply bending at the thecae junction. Arrowhead indicates protrusion of the plasma membrane (CFM, CCMP1335). Oogonia images are representative of 38 total images. **D:** Spermatogonium containing multiple spermatocytes seen as individual red (DNA stained) clusters (CFM, CCMP1335); representative of 8 images. **E:** Motile spermatocytes (in red, arrow) with moving flagella (arrowheads, CFM, CCMP1335, representative of 10 images). **F-G:** SEM images of spermatocytes (arrowhead) attached to early oogonia (SEM, CCMP1335, representative of 20 images). **H,I:** Auxospores; representative of 60 images in CCMP1015 (H, LM) and CCMP1335 (I, CFM) showing bulging where mother valve was attached (arrowhead). Two nuclei are visible in red following non-cytokinetic mitosis. **J:** Small parental cell (arrow) with initial cells produced by sexual reproduction to the left (partial valve view) and right (girdle view) indicated by the arrowheads (SEM, CCMP1335). **K:** 7 × 12 μm initial cell (LM); j and k representative of 12 images of CCMP1335. **L:** *T. weissflogii* auxospore (LM); representative of 12 similar images. **M:** *C. cryptica* spermatogonium (upper left) and vegetative cell (lower right). CFM shows stained DNA (red, arrow) and multiple nuclei in the spermatogonium. Arrowheads indicate chlorophyll autofluorescence (green). Oogonium (**N**, representative of 6 images) and auxospore (**O**, representative of 4 similar images) of *C. cryptica* (LM). Confocal microscopy images (b-e, i, m) show chlorophyll autofluorescence (green) and Hoescht 33342 stained DNA (red). Scale bars: **A**: 5 μm; **B-E**: 5 μm; **F**: 2 μm; **G**: 1 μm; **H-O**: 10 μm.

*T. pseudonana* spermatogonium harbored at least eight nuclei (Fig 3D), suggesting that a depauperating mitosis preceded meiosis [11, 35]. Sperm released were very small, about 1 μm, and flagellated (Figs 3E and S2E, S2F), but they often became entangled with other cells and debris [8, 36, 37]. Sperm cells attached to oogonia at the junction of the thecae for fertilization (Figs 3F, 3G and S2I, S2J) as shown in *T. punctigera* [38]. Also similar to *T. punctigera*, flagella were not visible at that stage, possibly because flagella are abandoned upon attachment to the oogonia [36].

Oogonia developed into auxospores and these conspicuous cell morphologies were always observed in cultures induced by ammonium. Auxospores were larger than vegetative cells and oogonia, ranging from about 6 to 20 μm in diameter, with most being 10-15 μm in diameter (Figs 3H, 3I and S2G, S2H, S2L). Auxospores were spherical, with most of the cellular contents localized to one side (Fig 3H, 3I and S2G, S2H, S2L) and sometimes showing slight distention where the mother valve was shed (Fig 3I), as described in *Stephanodiscus niagarae* [39]. Thecae remained attached in some cases, especially on smaller cells. Oogonia, auxospores, and spermatogonia in the other species studied displayed similar morphologies to those observed in *T. pseudonana* and other centric diatoms (Figs 3L-O, S2M-P and 4A) [11, 16, 28, 33, 35, 37, 38, 40, 41].

Changes in DNA content in *T. pseudonana* cells induced into sexuality by ammonium were observed using flow cytometry-based analysis. Fluorescence-activated cell sorting (FACS) analysis showed that as the culture progressed from exponential into stationary growth phases, the diploid population (Fig 4B) expanded to include DNA fluorescence intensities that were consistent with the presence of spermatogonangia and spermatogonia containing multiple gametes (Fig 4C). In late stationary phase, a new population was observed that had DNA fluorescence signals consistent with haploid sperm cells with little to no chlorophyll [13] (Fig 4D).

**Fig 4.**
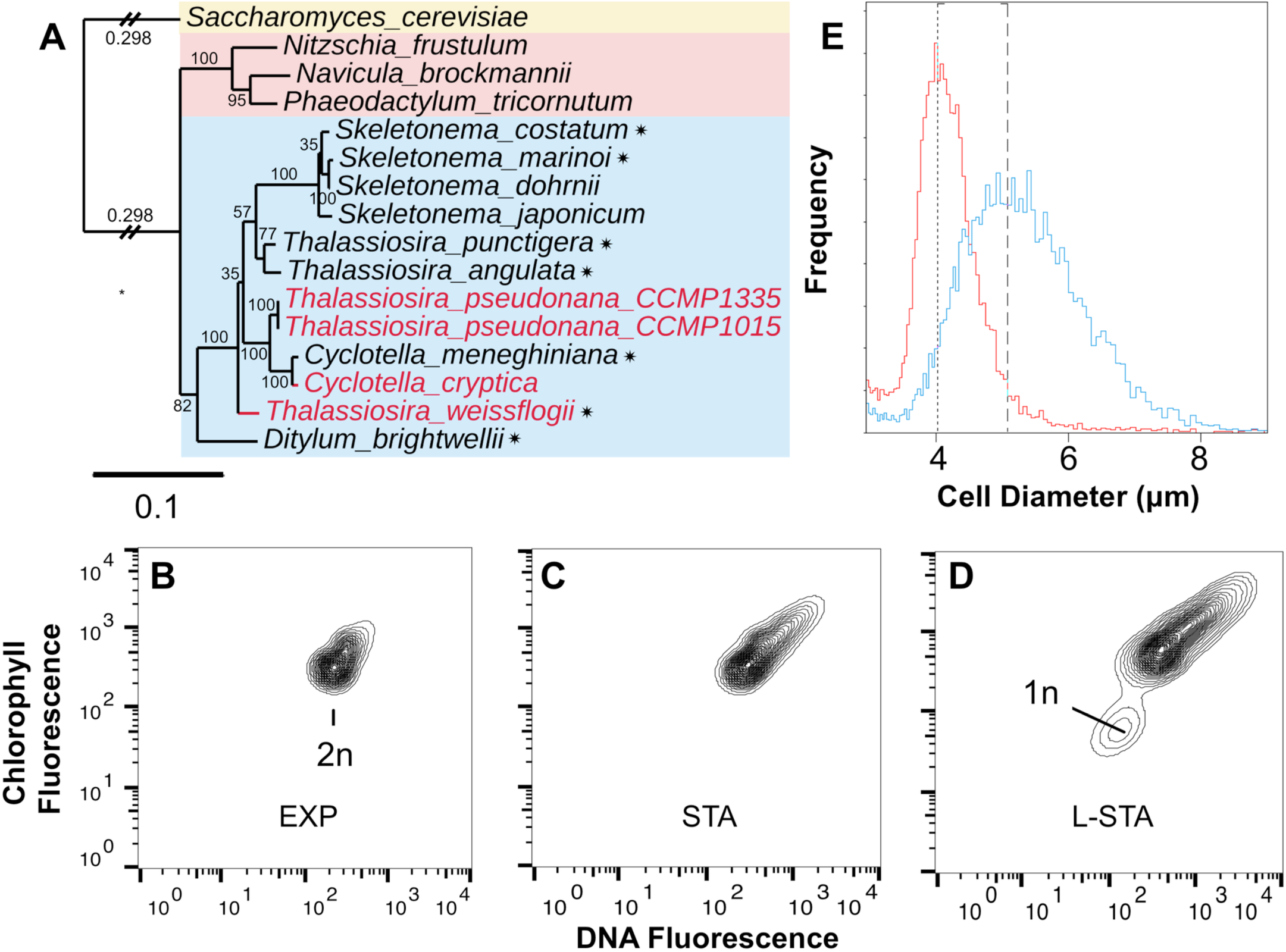
Evidence for meiosis and initial cells. **A:** 18s rRNA phylogeny of diatoms including pennates (pink rectangle), centrics (blue rectangle). Highlighted in red are the four strains induced into sexual reproduction in this study. Species for which some evidence already exists for sexual reproduction are starred [9, 13, 16, 28, 33, 38]. **B-D:** Changes in DNA and chlorophyll fluorescence in exponential (EXP), stationary (STA) and late stationary (L-STA) growth phases of *T. pseudonana* induced by ammonium; 30,000 events recorded, representative of two biological replicates. **E:** Coulter Counter distributions of cell diameter for *T. pseudonana* cultures in exponential phase of growth and maintained in NaNO_3_ (red) and after six successive 25% transfers to medium with ammonium (blue). Each new culture was allowed to remain in stationary phase for three days before the next 25% transfer was made. Single replicates of cultures with cell densities of 2.4 × 10^6^ ml^-1^ (NaNO_3_) and 2.3 × 10^6^ ml^-1^ (ammonium). Dashed lines are the mode for each peak.

Cell size restitution via auxosporulation produced progeny cells that were considerably larger than the parent cells from nitrate stock cultures. To induce a high proportion of the eligible cells into the sexual pathway we repeatedly propagated cultures in 800 μM ammonium with inoculum of 25%. This strategy raises the ratio between the exposure of cells to ammonium in stationary phase and the total number of cell divisions. Our findings can be explained by assuming that cells in nitrate stock cultures are already at or below the critical size threshold for induction into sexuality, but with each passage through growth and stationary phases in batch culture, only a fraction of the eligible cells are induced into the sexual cycle. The average cell diameters of the resulting cultures were larger relative to stock cultures maintained in nitrate (Fig 4E and S3). The *T. pseudonana* initial cells were 7-12 μm, the largest size reported for this species (Figs 3J, 3K, S2K). Presuming that cell size reduction during vegetative growth occurs in *T. pseudonana*, this process of cell size reduction and cell size restitution via ammonium induction have opposing influences on the average size of populations. These processes confound the ability to observe the impacts of sexual induction without experimental designs that maximize the percentage of the population induced into the sexual pathway and minimize the number of vegetative replications between episodes of induction.

We identified oogonia, male gametes, auxospores, and initial cells in cultures of the model centric diatom, *T. pseudonana* providing new evidence for sexuality in this species that was previously assumed to be asexual [10]. Although cell enlargement through asexual/apomictic mechanisms has been recorded in other species [42-44], the presence of all sexual cell types, and the expression of meiotic genes (discussed below), suggest apomixis is not the mechanism being used by *T. pseudonana* for cell enlargement. Furthermore, apomixis typically occurs in species that also undergo sexual reproduction [5]. Only spermatogenesis had previously been reported in *T. weissflogii* [13, 45], but we have now also documented induction of oogonia and auxospores by ammonium and subsequent formation of initial cells in this species. A major challenge in visualizing the morphological characteristics of these species is their smaller cell sizes compared to other species for which morphological details have been documented. Now that we have determined a reliable method for inducing the sexual morphologies, future studies will dissect additional details associated with the sexual pathways in these, and perhaps other species inducible by ammonium, to determine their variation from other centric diatoms. For example, the presence of auxospore scales, precise timing of fertilization and meiotic activity, repeated auxosporulation, and polyspermy events (e.g., [38]). The case of *T. pseudonana* also presents interesting questions about whether this species has retained the ability to reduce in cell size. It appears that *T. pseudonana* has the capacity to avoid clonal death by maintaining a relatively constant cell size (3-9 μM) [46-48]. Our experiments show that cells in this size range are inducible into the sexual pathway. Nevertheless, the question remains whether the progeny of induced small cells of *T. pseudonana* are capable of cell size reduction.

### Gene expression analysis of ammonium induced sexual morphologies

We used RNAseq to identify genes that were differentially expressed in conditions that triggered cell differentiation into sexual morphologies. We compared the transcriptomes of *T. pseudonana* harvested in exponential (EXP), stationary (STA), and late stationary phases (L-STA). Cells were grown in 100 μM NaNO_3_ or, to capture a dose-dependent change in gene expression, either 100 or 800 μM NH_4_Cl(S4 Fig). We identified genes that were significantly differentially expressed in multiple pairwise comparisons of growth phases and nitrogen sources (S1-S11 Tables). Next, we examined the statistical interactions of pairwise condition comparisons to identify genes with significantly greater or lesser magnitude changes in expression between growth stages in the presence of ammonium relative to 100 μM NaNO_3_ (Fig 5A and S5).

**Fig 5.**
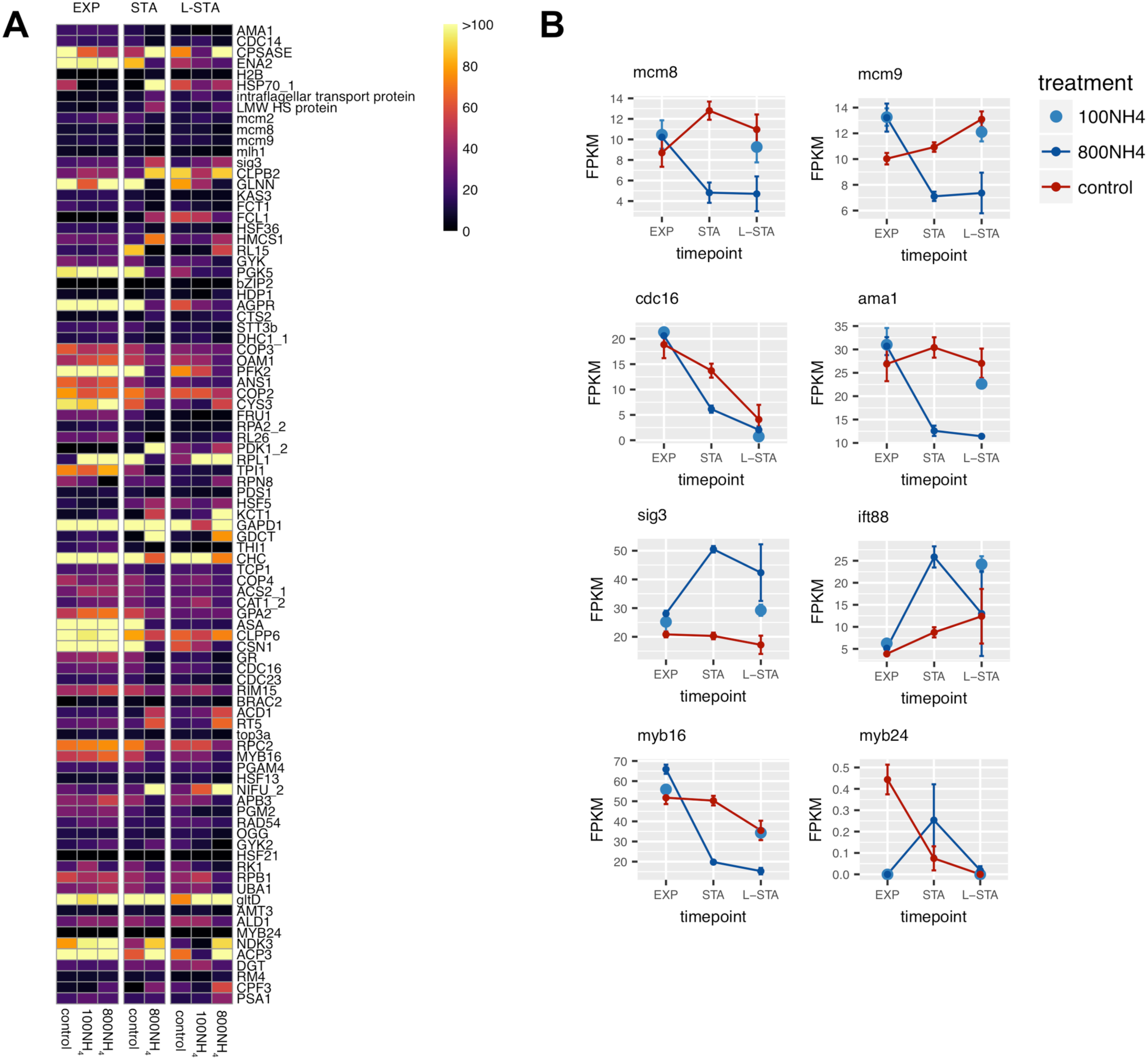
Transcriptomic evidence for sexual reproduction in *T. pseudonana*. **A:** Heat map of 89 genes having annotated functions that were differentially expressed during differentiation and sexual reproduction in *T. pseudonana* CCMP1335. Color indicates normalized expression value (FPKM) for each nitrogen treatment (control = 100 μM NO_3_^-^; 100NH_4_ = 100 μM NH>_4_^+^; 800NH_4_ = 800 μM NH_4_^+^) and growth phase (EXP, STA, L-STA). **B:** FPKM values of select genes across growth phases for each nitrogen treatment.

This conservative approach yielded a total of 1,274 genes in the four analyses of statistical interactions (S12-S15 Tables). A total of 89 of the genes have an annotated function (Fig. 5A;S16 Table). The set of 89 genes includes four meiotic genes (*mcm2, mcm8, mcm9, and mlh1*, Fig 4B and S6) that were also up-regulated in pennate diatoms during sexual reproduction [26]. The Mcm family of DNA helicases function in DNA replication, with *mcm8, mcm9*, and *mlh 1* having roles in double stranded break repair [49-51]. Mcm8 was one of the four genes related to meiosis that were upregulated in the pennate diatom, *Seminavis robusta* during treatment with a sex-inducing pheromone [17]. Eight genes in our list are homologous to yeast genes involved in meiosis [52] (Fig 5B, S17 Table), including genes that regulate the meiotic anaphase promoting complex (*cdc16*, *cdc23*, *ama1*) [53, 54] and *rad54*, a motor protein that regulates branch migration of Holliday junctions during homologous recombination [55]. Expression of genes encoding other RAD proteins (*rad50 and rad 51*) increased in pennate diatoms induced into meiosis [17, 26].

Of three ‘sexually induced genes’ that were up-regulated in *T. weissflogii* at the initiation of gametogenesis [45] and associated with sperm flagella mastigonemes [56], one, *sig3*, was significantly up-regulated in stationary phase compared to exponential phase (Fig 5B). In addition, a gene encoding an intraflagellar transport protein (IFT88) was also up-regulated in ammonium induced cells during stationary phase (Fig. 5B). An IFT system is required for flagellar assembly [57] and five genes encoding IFT particle proteins, including IFT88, and a kinesin-associated protein involved in anterograde transport were found in the *T. pseudonana* genome [58]. The genes encoding Sig3 and IFT88 are unique to flagellar structures, and their differential expression in ammonium induced *T. pseudonana* compared to the nitrate-grown control treatments provide additional evidence that ammonium induced spermatogenesis in this species.

The MYB factor and bZIP families of proteins are transcriptional regulators that control a variety of cell processes including stress responses, development, and differentiation in plants [59]. Expression of two genes having the characteristic R2R3 MYB DNA binding domains common in plants, *myb24 and myb16*, was generally lower in ammonium induced cultures compared to the nitrate grown controls (Fig 5B). In *Arabidopsis thaliana*, hormonal activation of *myb24* is required for stamen development and male fertility [60]. Whether *myb24, myb16*, and *bzip2* play roles in regulating gamete development or sex differentiation in diatoms remains to be determined.

The 1,274 genes provide new avenues to understand the evolution of sexuality in the Heterokont eukaryotic lineage. Diatoms emerged ~ 200 Mya, about 800 My after a eukaryotic heterotroph engulfed a red alga in the secondary endosymbiosis event that gave rise to the SAR eukaryotic supergroup [61]. Of 171 diatom genes of red algal origin [61], 17 were identified as differentially expressed in conditions that induced sexual reproduction (S18 Table). None of these genes are annotated in the *T. pseudonana* genome, but in red algae they are predicted to function in transport and plastid-targeted processes [23].

## Conclusions

That some of the most well studied centric diatoms were never observed undergoing sexual reproduction was a mystery. Possibly even more elusive was the ability to reliably control or induce the sexual pathway of centric diatoms in the laboratory [ 10] despite a myriad of efforts that ranged from sweet-talk to torture. Factors that have limited progress in this field center on the problem that even under ‘favorable environmental conditions’ that result in the sexual lifecycle, only a fraction of cells undergo sexual reproduction (Fig 2). Thus, capturing the sexual event requires near constant visual observation because (a) only cells that have become sufficiently small and reach the critical size threshold can undergo sexual reproduction [62], (b) only a fraction of those size-eligible cells may undergo sexual reproduction [8, 15, 16, 31], (c) there has been a lack of understanding about what constitutes conditions that are ‘favorable’ for triggering diatom sex [5, 7], and (d) morphological changes indicative of sex may not be recognized by untrained scientists [63].

Our results provide strong evidence that *T. pseudonana* is a sexual organism, expressing the major morphologies associated with the sexual pathway that result in enlarged initial cells. Furthermore, the sexual pathway was reliably induced in *T. pseudonana*, and two other centric diatom species by exposure of size-eligible cells to ammonium. Ammonium triggered formation of sexual cells in a dose-dependent manner and significant changes in expression of genes involved in meiosis, spermatocyte flagellar structures and assembly, and sex differentiation. RNAseq analysis revealed many more genes with unknown functions that were expressed under conditions of sexual differentiation. Other genes involved in sex are likely to have been missed by our analysis because their changes in expression were masked by the mixed population of asexual and sexual cells, or they were not captured in the coarse time-resolution of sampling used in this study. Nevertheless, our discoveries resolve two persistent mysteries that have plagued diatom researchers. Furthermore, the RNAseq data provide a subset of genes that can be used to study the molecular ecology of diatoms.

The ecology of centric diatom sexual reproduction that can be inferred from our findings appears best described as synchronous sexuality [12] triggered by ammonium in cells experiencing growth stress. Asexual cell cycle arrest appears to be prerequisite to activation of the diatom sexual life cycle [13, 14, 28, 29]. In the environment, diatoms bloom following elevated nutrient concentrations driven by vertical mixing, coastal upwelling, or river inputs and the bloom reaches its peak biomass when essential nutrients are depleted. Within a week, the bulk of a bloom can be consumed by heterotrophic protists [64] that excrete ammonium to maintain homeostatic elemental composition [65]. We propose that ammonium released by grazers at bloom climax may be a principal ecological trigger for sexual morphologies in centric diatoms. Synchronization of sexuality at the onset of resource depletion (stationary phase) increases the chances for successful fertilization because cell density is at its maximum [12]. Environmental concentrations of ammonium in the environment rarely reach the concentrations used in this study to demonstrate the dose response effect on sexuality. Other methods that have sometimes successfully triggered sexual reproduction in other species are similarly unusual compared to environmental conditions. For example, the magnitude of the salinity shifts used to induce sexual reproduction in *Skeletonema marioni* in the laboratory do not occur in the Baltic sea [16]. Nevertheless, pulses of ammonium, shifts in salinity, and other environmental fluctuations do occur in aquatic ecosystems, and provided the other conditions for sexuality are met (e.g., cell size threshold, stress, population density), are likely to induce sexuality in at least a small fraction of a population. The presence of ammonium and the onset of stationary phase also point to involvement of another growth factor whose depletion triggers sexual reproduction. The specific collection of factors that lead to sexual reproduction in diatoms in the environment is not yet known and neither is whether ammonium is a direct or indirect trigger of sexuality [4]. Nevertheless, this work suggests an intriguing ecological role for ammonium in the mechanisms underlying sexuality in centric diatoms and will certainly be a valuable tool to control sexuality in the laboratory.

The identification of ammonium as a reliable inducer of sexuality in *T. pseudonana* and other centric diatoms has the potential to shift perspectives on diatom ecology, open avenues for the experimental investigation of diatom reproductive mechanisms, and provide tools for genetic manipulation of centric diatoms that have not heretofore been available. Diatom blooms have a global impact but the factors that control these blooms and their demise are complex and a consensus has not been reached about these processes. Our evidence suggests that induction of sexuality may play a vital role in diatom bloom conclusion and the production of genetic diversity that seeds future blooms [66]. Our analysis suggests an involvement of genes of red-algal origin, providing new lines of evolutionary enquiry. Interest in diatoms for biotechnological applications is high due to their uses in biofuels, materials chemistry and medicine. Our work will likely propel this exploration by enabling improved breeding and genetic modification to control and understand unique diatom traits.

## Materials and methods

Stock cultures of *T. pseudonana* (CCMP1335) were maintained in f/2 medium [67] with 200 μM NaNO_3_ under continuous sub-saturating light at 18°C. Sexual cells were quantified in triplicate cultures of *T. pseudonana* (CCMP1335 and CCMP1015)*, T. weissflogii* (CCMP1336) and *C. cryptica* (CCMP332) (all obtained from NCMA) grown in f/2 amended with NaNO_3_ or NH_4_Cl and grown at 18°C under 50 μE continuous light, or under variable light intensities/cycles as shown in Table 1. Cell populations were quantified using a Coulter counter (Beckman-Coulter, Indianapolis, Indiana). Oogonia and auxospores were counted using a hemocytometer.

We found that a modified f/2 medium yielded better cell images using light and confocal microscopy. This medium contained 0.939 mM KCl, 0.802 mM NO_3_^-^, 1 mM NH_4_Cl, 0.05 mM glycine, 0.01 mM methionine, 0.078 mM pyruvate, 0.84 μM pantothenate, 0.985 μM 4-amino-5- hydroxymethyl-2-methylpyrimidine, 0.3 μM thiamine, 0.002 μM biotin, 0.117 μM FeCl_3_*6H_2_O, 0.009 μM MnCl2*4H_2_O, 0.0008 μM ZnSO_4_*7H_2_O, 0.0005 μM CoCl_2_*6H_2_O, 0.0003 μM Na_2_MoO_4_*2H_2_O, 0.001 μM Na_2_SeO_3_, and 0.001 μM NiCl_2_*6H_2_O, and sparged with filter-sterilized carbon dioxide and air for 8 hours and overnight respectively. To view DNA, 1 ml live samples were stained with 5 μl 1.62 μM Hoescht 33342 (0.2 μm filtered) for 10 min. For scanning electron microscopy (SEM), 1 ml samples were diluted 1:3 with sterile f/2 and syringe filtered onto 13 mm 0.2 μm polycarbonate filters using a Swinnex filter unit. The filter was washed with 4 ml f/2 containing 0.5% gluteraldehyde and left submerged for at least 24 hours, followed by a series of 4 ml washes: 0.2 μm filtered 80%, 60%, 40%, 20% and 0% f/2, followed by 20%, 40%, 60% 80% and 100% ethanol, before critical point drying. SEM imaging was done at the Oregon State University Electron Microscope facility.

For flow cytometry 1 ml culture samples were fixed with 1 μl gluteraldehyde (50%) and stained with 10 μl Sybr green mix (1:25 dilution Sybr green in 0.01M Tris-EDTA, pH 8.0) for 30 min. Samples were run on a FACScan flow cytometer (Becton Dickinson, Franklin Lakes, New Jersey). Settings were FL1=582 and FL3=450 for unstained cells and FL1=450 and FL3=450 for stained cells.

For RNAseq analysis, 1.61×10^8^ − 1.10×10^9^ cells from triplicate independent cultures were filtered onto 0.8 μm 47 mm polycarbonate filters during exponential, stationary and late stationary phases and flash frozen in liquid nitrogen. RNA was extracted using a Qiagen RNeasy midi kit according to modified manufacturer’s instructions [68]. Silica beads (0.5 mm) were added to the cells and lysis buffer and vortexed until homogeneous before being filtered through Qiashredder columns to remove large particles. Eluted RNA was subjected to off column RNase free DNase I treatment and secondary purification according to manufacturer’s recommendations. Total RNA was prepared and sequenced as a 150 bp single end library on an Illumina HiSeq 3000 at the Center for Genome Research and Biocomputing at Oregon State University. Sequencing data/interaction analyses were conducted using the Ballgown pipeline [69]. Sequencing reads were trimmed to remove sequencing adapters using BBDuk v. with the parameters “ktrim=r k=23 mink=9 hdist=1 minlength=100 tpe tbo” [70]. Reads aligned to the *T. pseudonana* reference genome (NCBI accession GCA_000149405.2) using HISAT2 v. 2.0.4 with the parameters “--min-intronlen 20 --max-intronlen 1500 --rna-strandness F --dta-cufflinks” [71]. Transcripts were assembled for each dataset and merged using Stringtie v 1.2.4 [72]. Pairwise differential expression analyses for genes were performed using the “stattest” function in Ballgown version 2.2.0 [73]. Interaction effects were tested by comparing the models with (timepoint + treatment + timepoint * treatment) and without (timepoint + treatment) the interaction term using the custom model option in the “stattest” function.

For construction of the phylogenetic tree, 18s rRNA sequences were obtained from the Silva database and aligned using Muscle v3.8.31 (default settings) [74]. A genome editor (BioEdit) was used to manually trim off overhanging sequence. The tree was built using RAxML-HPC v8.0.26 using the GTRCAT model, “-f a” option, and 1000 bootstrap replicates [75]. A visual representation was created using the TreeDyn [76] tool through LIRMM (phylogeny.fr) [77].

All RNAseq data have been deposited to NCBI under BioProject ID PRJNA391000.

## Acknowledgements

The authors thank Kelsey McBeain for assistance with confocal microscopy imaging and the Electron Microscope Facility at Oregon State University for scanning electron micrographs.

## Supporting Information

**S1 Fig. Ammonium induces sexual morphologies in *T. weissflogii* (A) and *C. cryptica* (B).** Proportion of sexual cells (oogonia and auxospores) relative to the total population in cultures supplemented with NH_4_Cl or NaNO_3_. An average of 120 and 107 cells were counted per replicate of *T. weissflogii* and *C. cryptica*, respectively, throughout the growth curve, but oogonia and auxospores were only observed beginning in stationary phase; independent cultures n=3, data are mean values, error bars are s.d.. Inset: corresponding growth curve linking the onset of stationary phase with first appearance of sexual cells on day 10 (**A**) and 17 (**B**).

**S2 Fig. The different life stages in *T. pseudonana* (A-L)**, ***T. weissflogii* (M,N) and *C. cryptica* (O,P)**. **A:** SEM of vegetative cells (CCMP1335). **B-D:** SEM (B) and CFM images of CCMP1335 oogonia, displaying separation of the thecae and expansion of the membrane. **E:** CFM image of flagellated spermatocytes with stained DNA (arrowheads), **F, G**. Epifluorescence (F) and LM images of the same view. In F, an active, flagellated spermatocyte (arrowhead) possibly associated with an auxospore surface is revealed by lateral light from fluorescence of DNA (blue) and chlorophyll (red). **H,L:** Auxospores of CCMP1015 and CCMP1335 respectively (CFM). **I,J:** Individual spermatocytes attached to oogonia (SEM). **K:** Initial cells of *T. pseudonana* CCMP1335 (LM). **M,N:** *T. weissflogii* vegetative cells (M; LM) and auxospore (N; LM). **O,P:** *C. cryptica* oogonia (O; LM) and auxospores (P; LM). CFM images (C-E, H, L) show fluorescence of DNA in red and chlorophyll in green.

**S3 Fig.** Coulter Counter distributions of cell diameter for *T. weissflogii* (**A**) and *C. cryptica* (**B**) cultures in exponential phases of growth and maintained in NaNO_3_ (red) and after two successive 25% transfers to media with ammonium (blue), Each new culture was allowed to remain in stationary phase for three days before the next 25% transfer was made. Single replicates. Dashed lines are the mode for each peak. Cell densities in (**A**) are 2.2 × 10^5^ ml^-1^ (NaNO_3_) and 3.3 × 10^5^ ml^-1^ (ammonium) 1.6 × 10 ml^-1^ 2.2 × 10^6^ and (**B**) are 1.6 × 10^6^ ml^-1^(NaNO_3_) and 2.2 × 10^6^ml^-1^ (ammonium).

**S4 Fig. *T. pseudonana* CCMP1335 growth and collection for RNAseq analysis.** Three independent cultures of each nitrogen treatment were harvested 3, 5, and 8 days after inoculation (down arrows) in exponential (EXP), stationary (STA) and late stationary phases (L-STA). The 100uM NH_4_^+^ STA treatment did not yield sufficient RNA for analysis.

**S5 Fig. Interaction analysis workflow of RNAseq data.** Growth phase A vs. B is EXP vs. STA, EXP vs. L-STA, or STA vs. L-STA, respectively. Δexp is the magnitude of change in gene expression between growth phases for the different nitrogen treatments.

**S6 Fig. Expression values (FPKM) of 15 selected genes across the three growth phases for each nitrogen treatment.**

**Table S1. Pairwise comparison, Stationary phase 800 μM Ammonium vs. exponential phase 800 μM Ammonium.**

**Table S2. Pairwise comparison, Late-stationary phase 800 μM Ammonium vs. exponential phase 800 μM Ammonium.**

**Table S3. Pairwise comparison, Exponential phase 100 μM Ammonium vs. Late stationary phase 100 μM Ammonium.**

**Table S4. Pairwise comparison, Stationary phase 800 μM Ammonium vs. Late stationary phase 800 μM Ammonium.**

**Table S5. Pairwise comparison, Exponential phase nitrate vs. Exponential phase 800 μM Ammonium.**

**Table S6. Pairwise comparison, Stationary phase nitrate vs. Stationary phase 800 μM Ammonium.**

**Table S7. Pairwise comparison, Late Stationary phase nitrate vs. Late Stationary phase 800 μM Ammonium.**

**Table S8. Pairwise comparison, Exponential phase nitrate vs. Exponential phase 100 μM Ammonium.**

**Table S9. Pairwise comparison, Late Stationary phase nitrate vs. Late Stationary phase 100 μM Ammonium.**

**Table S10. Pairwise comparison, Exponential phase 100 μM Ammonium vs. Exponential phase 800 μM Ammonium.**

**Table S11. Pairwise comparison, Late Stationary phase 100 μM Ammonium vs. Late Stationary phase 800 μM Ammonium.**

**Table S12. Interaction analysis, Exponential - Stationary phases (800 μM Ammonium v. Nitrate control).**

**Table S13. Interaction analysis, Exponential - Late Stationary phases (800 μM Ammonium v. Nitrate control).**

**Table S14. Interaction analysis, Exponential - Late Stationary phases (100 μM Ammonium v. Nitrate control).**

**Table S15. Interaction analysis, Stationary - Late Stationary phases (800 μM Ammonium v. Nitrate control).**

**Table S16. Genes that were identified as differentially expressed in conditions that triggered cell differentiation and sexual reproduction and have annotated functions.**

**Table S17. Genes that were identified as differentially expressed in conditions that triggered cell differentiation and sexual reproduction and are homologous to yeast genes involved in meiosis.**

**Table S18. Genes that were identified as differentially expressed in conditions that triggered cell differentiation and sexual reproduction and are of red algal origin.**

